# Human milk contains a heterogeneous population of EVs and microRNAs that resist simulated digestion

**DOI:** 10.64898/2026.02.10.705086

**Authors:** Zeinab Husseini, Elena C. Pulido-Mateos

## Abstract

Mother’s milk is known for its crucial roles in infant health and development. Probably the most commonly studied effects of milk are those on an infant’s intestinal barrier, metabolism, and immunity. While these functions of milk were mostly attributed to its protein and fat content, recent evidence points to a potential role of milk microRNAs in these processes. MicroRNAs are small non-coding RNAs that can fine-tune gene expression at the post-transcriptional level. Human milk (HM) is rich in microRNAs, which are mainly found associated with milk extracellular vesicles (EVs). HM microRNAs are proposed to transfer from mother to infant via breastfeeding and execute gene regulatory functions in infant cells. For microRNAs to be able to act as “genetic programmers” rather than mere nutritional molecules, they should resist digestion in the infant’s gastrointestinal tract. Milk EVs are believed to protect microRNAs against degradation and facilitate their delivery to the cells. Here, we used two lots of pasteurized HM that were originally destined for human milk banks. We showed that HM contains different populations of EVs with different physicochemical properties, similar to those previously identified in commercial bovine milk. We also showed that these EVs, which are often discarded, contain the majority of HM microRNAs. Finally, we showed that three highly abundant milk microRNAs resisted differentially to infant’s simulated digestion conditions, with a relatively small number of microRNAs surviving a two-hour digestion. Milk microRNA copy numbers surviving digestion may be too low to influence gene expression in infant cells.

**Highlights:**

- Pasteurized HM contains heterogeneous populations of EVs.
- These EVs associate with the majority of HM microRNAs.
- Different microRNAs show varying stability during infant digestion.
- The copy number of milk miR-148a-3p surviving digestion might be too low to influence gene expression in infant cells.

## Introduction

Human milk (HM) represents a complex pool of nutritional and bioactive molecules that confer health and developmental benefits for the infant ^(1)^. The beneficial effects of HM on infants’ biological systems are well-documented with implications that extend across both short- and long-term runs. Notably, HM promotes the development and maturation of the infant’s intestinal barrier, immune system, and central nervous system, and regulates early metabolic processes ^(2–4)^. While nutritional components in HM play a fundamental role, many of its health-promoting effects are attributed to bioactive molecules with beyond-nutritional values ^(5)^. These include growth factors (insulin, IGF-I, IGF-II, EGF), antibodies, antimicrobial proteins, as well as maternal immune cells, which impart the infant with passive immunity ^(6,7)^.

In addition to these well-characterized bioactive molecules, HM also contains extracellular vesicles (EVs) and microRNAs that are increasingly recognized for their potential roles in immunity, metabolism, and development ^(8–15)^. New EV populations were unveiled and characterized in different cow milk samples ^(16–19)^. A study on pasteurized commercial cow milk identified four EV populations (pelleted at 12,000 × g (12K), 35,000 × g (35K), 70,000 × g (70K), and 100,000 × g (100K)) and showed that the majority of commercial cow milk microRNAs are associated with these EVs ^(18,19)^. Milk EVs are believed to associate with microRNAs, protect them against degradation during digestion, and deliver them to the infant’s cells, where they could function as gene regulatory molecules ^(20–24)^.

Since the survival of milk microRNAs against digestion is a prerequisite for their bioactivity in the consumer, resistance of milk microRNAs to digestion has been investigated in different contexts and using different milk samples. For instance, a fraction of microRNAs in pasteurized commercial cow milk was shown to resist *in vitro* digestion, simulating that of an adult ^(25)^. Since the gastrointestinal (GI) tract digestion conditions are different between adults and infants, the resistance of HM microRNAs to digestion should be assessed using conditions more similar to those of an infant. Compared to adults, infants have milder digestion (higher gastric pH, lower enzymes) and a more permeable gut, suggesting higher survivability, transcytosis, and absorption of dietary molecules, including milk microRNAs ^(26–31)^. Two studies assessed the stability of microRNAs from HM against infant simulated digestion, albeit using a less complex approach compared to the one used on commercial cow milk ^(32,33)^.

Notably, most insights into HM microRNA stability have been drawn from raw, non-processed milk. Nonetheless, when breastfeeding is not possible, infants are often provided formula or HM from milk banks. HM banks routinely use pasteurization to ensure microbiological safety. Pasteurization can affect the yield of milk microRNAs as well as the integrity and count of milk EVs ^(34–37)^. Milk pasteurization could thus influence the number of microRNAs potentially surviving after digestion and hence affect their bioactivity in infant cells.

Building on previous findings, we sought to assess whether pasteurized HM retains microRNA integrity under infant-like digestion conditions using the TNO gastro-intestinal model (TIM-1), which dynamically simulates the four compartments of the GI tract ^(27,38)^. Additionally, we aimed to investigate whether the newly identified EV populations reported in cow milk are also present in HM. For that, we used the differential ultracentrifugation pipeline for EV isolation from ^(39)^ to isolate the four EV populations (12K, 35K, 70K, and 100K). Since these EV populations were proposed to protect microRNAs from digestion, we assessed whether microRNAs in HM are associated with these EV populations and hence might be contributing to their stability.

## Materials and Methods

### Milk samples

This study was conducted according to the guidelines laid down in the Declaration of Helsinki, and all procedures involving human subjects were approved by the Comités d’éthique de la recherche avec des êtres humains de l’Université Laval under the project number 2020-4843. Milk samples were obtained under a Material Transfer Agreement between Héma Québec and Université Laval, upon request by Dr. Patrick Provost. Human milk samples used in this study were obtained from the Public Mothers’ Milk Bank at Héma-Québec center, Quebec City, QC, Canada. Each milk lot (1 liter) represents a pool of samples collected from 36 donor mothers (see **Supplementary Fig.S1**). These lots were pasteurized (63 °C for 30 min) and then tested for sterility. As they failed to pass the sterility test, they were deemed not adequate to be consumed by infants and were thus donated for research. The samples were collected from Héma-Québec on dry ice and were directly transferred (while still frozen) to storage at -80 °C until further analysis and experimentation.

### TNO TIM-1 in vitro digestion system

TNO TIM-1 is a computer-controlled system composed of 4 main digestive compartments mimicking the stomach, duodenum, jejunum, and ileum, and is connected through peristaltic valve pumps ^(38)^. Each compartment is made up of a glass jacket surrounding a flexible silicone membrane, which compresses and relaxes to allow for the peristaltic mixing of the chyme. The pH, heat, and other parameters are continuously monitored through computer-connected sensors. Accordingly, digestive enzymes (e.g., trypsin, pepsin, lipase) and reagents (e.g., HCl and bicarbonate) are injected into each compartment to maintain physiologically relevant conditions.

### TIM-1 in vitro digestion

Human milk (100 ml) was introduced into TIM-1 and was digested for 2 h at 37 °C. To simulate the digestion of a liquid sample in a healthy human infant aged 1 to 6 months, parameters were adapted based on data compared across multiple literature resources ^(26,28,40,41)^ (Detailed digestion parameters are presented in **Table 1**). Gastric pH was initially set at 6.5 and was adjusted to gradually decrease over time, to reach 6.3 at 40 min, 6.0 at 60 min, 5.0 at 90 min, and 4.5 at 120 min. The intestinal pH values (6.5 in the duodenum, 6.8 in the jejunum, and 7.2 in the ileum) were set according to pediatric gastro-intestinal conditions ^(42)^, which are similar to those reported for adults. Enzymes used for digestion included pepsin and gastric lipase, which were added at 75% of the adult levels. Additionally, intestinal enzymes such as trypsin, amylase, lipase, ribonuclease, and proteases (added in the form of pancreatin) were also used at 75% of the levels typically used for adults. Gastric half-emptying time (t / = 60 min) and β = 1.5 were applied to regulate gastric transit ^(42)^. Jejunal and ileal compartments were coupled to semi-permeable hollow-fiber dialysis units (SUREFLUX-07L, Nipro Corporation, Settsu, Osaka, Japan) to simulate small-intestinal absorption.

**Table 1.** TIM-1 digestion parameters. The digestion parameters were adjusted to simulate the digestion of a liquid sample in a healthy infant. The parameters were continuously computer-monitored to determine the required volume of secretions per compartment. * Gastric and ileal deliveries are modelled with a power exponential formula: f = 1–2-(t/t1/2) β, where f represents the fraction of the meal delivered, t the time of delivery, t1/2 the half-time of delivery, and β a coefficient describing the shape of the curve.

| Compartment | Initial content and secretions | Volume (ml) | pH | $t_{1/2}$ and $\beta$ |
| --- | --- | --- | --- | --- |
| Stomach | Initial content: 3 mL of Gastric residue = 3ml with 6750 U pepsin, 280 U lipase per litre, and simulated gastric fluid: $\text{Na}^+$ 94 mmol/L, $\text{K}^+$ 13.2 mmol/L, $\text{Cl}^-$ 107.2 mmol/L.<br>pH 3.5<br>Secretions:<br>Gastric juice; pH 6 – 0.25 ml/min<br>675 U pepsin, 28 U lipase /mL.<br>Simulated gastric fluid: $\text{Na}^+$ 94mmol/L, $\text{K}^+$ 13.2mmol/L, $\text{Cl}^-$ 107.2 mmol/L.<br>Water or HCl 0.5 M – 0.25 ml/min | 125 | 0/6.5<br>40/6.3<br>60/6<br>90/5<br>120/4.5<br>150/3.8<br>180/3.5 | 60 and 1.5 |
| Duodenum | Initial content: 15 g of a 6 % pancreatin solution (8xUSP), 30 g of a 6 % porcine bile solution, 500ul of 2 mg/mL trypsin solution and 15 g of small intestine electrolyte solution (SIES: NaCl 5 g/L, KCl 0.6 g/L, CaCl<sub>2</sub> 0.25 g/L)<br>Secretions:<br>6 % pancreatin solution (8xUSP) – 0.25 mL/min<br>6 % bile solution (first 30 min), then 3 % – 0.50 mL/min<br>SIES or NaHCO<sub>3</sub> 0.5 M – 0.5 mL/min | 55 | 6.5 | <150 and 1.8 |
| **Jejunum** | Initial content: SIES<br>Secretions: NaHCO<sub>3</sub> 0.5 M if necessary<br>Dialysis fluid : SIES – 10 mL/min | 115 | 6.8 |  |
| **Ileum** | Initial content: SIES<br>Secretions: NaHCO<sub>3</sub> 0.5 M if necessary<br>Dialysis fluid : SIES – 10 mL/min | 115 | 7.2 |  |

### TIM-1 Media

Freshly prepared working solutions contained pepsin derived from porcine gastric mucosa (Sigma-Aldrich, P6887), trypsin sourced from porcine pancreas (Sigma-Aldrich, St. Louis, MO, USA, T9201), pancreatin extracted from porcine pancreas (Sigma-Aldrich, St. Louis, MO, USA, 8 × USP, P7545), and lipase from *Rhizopus oryzae* (Amano Enzyme USA Co., Ltd., Elgin, IL, USA, LIPASE DF-DS), all dissolved in sterile deionized water. To mimic bile secretion, porcine bile (Sigma-Aldrich, St. Louis, MO, USA, B8631) was added at 75 % of an adult dose, corresponding to an initial duodenal bile salt concentration of 7.5 to 11.25 mM, which decreases progressively in the TIM-1 to approximately 3.75 mM after 2 h of digestion ^(43)^. The electrolyte solutions consisted of a gastric electrolyte mixture (Na+ 94 mmol/L, K+ 13.2 mmol/L, Cl^-^ 107.2 mmol/L) and a small intestinal electrolyte solution (NaCl 5 g/L, KCl 0.6 g/L, CaCl2 0.25 g/L).

### TIM-1 sample collection

Samples (1 ml) were collected from each of the four TIM-1 compartments at 0, 30, 60, and 120 min. Also, at each time point, the effluent was collected, and its volume was recorded. Samples were flash-frozen on dry ice and stored at −80 °C for later RT-qPCR analysis.

### Isolation of human milk extracellular vesicles by differential ultracentrifugation

Milk EVs were obtained by following a previously described protocol, with slight modifications, that allows quick isolation of milk EVs with very little contaminating proteins ^(39)^. Milk was centrifuged at 4,500 × g for 30 min to remove potentially remaining cells or cell debris (pellet) and fat layer (upper floating layer). The partially skim milk obtained (middle layer) was then subjected to another centrifugation at 4, 500 × g for another 30 min at 4 °C to remove the remaining fat layer. The skim milk obtained was filtered through 0.45 µm filter and was then added to 2% sodium citrate at 1:1 ratio and kept mixing for 20 min at 4 °C. The Milk: sodium citrate mixture was subjected to successive differential ultracentrifugation steps at 12,000 × g (12K) for 2 h, 35,000 × g (35K) for 1 h, then 70,000 × g (70K) for 1 h, and 100,000 × g (100K) for 1 h at 4 °C in a Sorvall WX TL-100 ultracentrifuge, equipped with a T-1250 rotor (both Thermo Fisher Scientific, Waltham, MA, USA). After each step, the pellets were suspended in 1 mL of 0.22 µm filtered sterile phosphate-buffered saline (PBS) pH 7.4.

### Dynamic light scattering

The four pellets obtained upon ultracentrifugation were diluted 10X or 100X in 0.22 µm-filtered PBS and deposited in an ultraviolet cuvette (VWR, Radnor, PA, USA, Cat. No. 47743-834). The hydrodynamic size, derived count rate, and polydispersity index were measured at 37 °C with three measurements for each using the Zetasizer Nano-ZS dynamic light scattering (DLS) measurement system (Malvern, Worcestershire, UK).

### Protein dosage

Protein dosage for all samples was done using the Pierce BCA Protein Assay Kit (Thermo Fisher Scientific, Waltham, MA, USA, Cat. No. 23227) as per manufacturer’s instructions. Absorbances were read at 562 □ nm using an EPOCH2 microplate reader (BioTek Instruments, Winooski, VT, USA).

### Western Blot

Proteins isolated from EVs were mixed with 6X loading buffer (0.15M Tris pH 6.8, 1.2% SDS, 30% glycerol, 1.8% bromophenol blue), heated at 95 °C for 10 min, and subjected to 10% or 12.5% sodium dodecyl sulfate (SDS)-polyacrylamide gel electrophoresis. Proteins were transferred to 0.45 μm (Millipore, Burlington, MA, USA, Cat. No. IPVH85R) pore size polyvinylidene fluoride membranes. The membranes were blocked with fat-free milk solution (5% milk powder in Tris buffer saline (TBS) with 0.1% Tween 20) for 1 h at room temperature. Membranes were then incubated with primary antibody overnight at 4 °C with monoclonal anti-TSG101 (Abcam, Cambridge, UK, clone 4A10, Cat. No. ab83, diluted 1/500), monoclonal anti-HSP70 (Becton Dickinson (BD), Franklin Lakes, NJ, USA, Cat. No. 554243, diluted 1/500), monoclonal anti-ALIX (Santa Cruz Biotechnology, Dallas, TX, USA, clone 3A9, Cat. No. sc-53,538, 1/500), polyclonal anti-CD63 (Santa Cruz Biotechnology, Dallas, TX, USA, Cat. No. sc15363, diluted, 1/1,000), and monoclonal anti-XDH (Santa Cruz Biotechnology, Dallas, TX, USA, Cat. No. sc-20991, diluted 1/1,000). Primary antibodies were diluted in 5% fat-free milk. Membranes were then washed three times with TBS tween (3 × 10 min), incubated for 1 h with horseradish peroxidase (HRP)-conjugated secondary anti-mouse (PerkinElmer, Waltham, MA, USA, Cat. No. NEF812001EA, diluted 1/10,000) or anti-rabbit (PerkinElmer, Waltham, MA, USA, Cat. No. NEF822001EA, diluted 1/15,000). Secondary antibodies were diluted in 5% fat-free milk. Membranes were then washed three times in TBST (3 × 10 min). Western blot signals were revealed with clarity western enhanced chemiluminescence (Bio-Rad Laboratories, Hercules, CA, USA, Cat. No.1705061). The chemiluminescent signal was captured by exposure to the HyBlot CL® Autoradiography 8 x 10”, 100 films (Thomas Scientific, Swedesboro, NJ, USA, Cat. No. 1141J52) and developed using the Konica SRX-101A X-ray film processor (Konica Minolta, Tokyo, Japan).

### RNA extraction from milk samples (skim milk, EVs and supernatant) and TIM-1 digests

RNA extraction was performed as per manufacturer’s instructions with slight modifications. Briefly, 1 ml of RNAzol RT (Sigma-Aldrich, St. Louis, MO, USA, Cat. No. R4533) spiked with 0.15 μL UniSp2 exogenous RNA (QIAGEN, Hilden, NW, Germany, Cat. No. 339390) and added to 250 μL skim milk, milk supernatant (SN), milk EVs, or samples collected from TIM-1. We also extracted RNA from samples collected from TIM-1 at T0 before the addition of milk. UniSp2 was used as an exogenous RNA to monitor for variations during RNA extraction. The above mix was added to 400 μL nuclease-free water and mixed vigorously by shaking for 20 s and then incubated at room temperature for 15 min. Next, the mix was centrifuged at 12,000 × g for 15 min at 4°C, and the supernatant (1,200 μL) was recovered. GlycoBlue (Thermo Fisher Scientific, Waltham, MA, USA, Cat. No. AM9516) and an equal volume of isopropanol were added, mixed gently, incubated for 10 min at room temperature, and centrifuged at 12,000 × g for 10 min at 4 °C. The supernatant was carefully removed, and the pellet was washed twice with 500 µL of 75% ethanol, centrifuging each time at 12,000 × g for 5 min at 4 °C. After air-drying for 10 min, the pellet was resuspended in nuclease-free water and RNA was treated with DNase-I (Invitrogen, Carlsbad, CA, USA, DNA-free™ DNA Removal Kit Cat. No. AM1906) in accordance with the manufacturer’s protocol.

### RT-qPCR of milk samples (EVs, skim milk, and supernatant) and TIM-1 digests

For cDNA generation, miRCURY LNA RT Kit (QIAGEN, Hilden, NW, Germany, Cat. No. 339340) was used. UniSp6 RNA, included in the miRCURY LNA RT Kit, was spiked in all RT reactions to control for cDNA synthesis and PCR efficiency. Next, cDNA was diluted 10X and qPCR was performed using miRCURY LNA SYBR Green PCR Kits (QIAGEN, Hilden, NW, Germany, Cat. No. 339345) in 0.1 mL MicroAmp Fast Optical 96-Well Reaction Plate (Applied Biosystem, Foster City, CA, USA, Cat. No. 4346907) in a StepOne Real-Time PCR instrument (Thermo Fisher Scientific, Waltham, MA, USA) and the LNA PCR primer assays for UniSp2 (QIAGEN, Hilden, NW, Germany, Cat. No. 339306, GeneGlobe ID. YP00203950), UniSp6 (QIAGEN, Hilden, NW, Germany, Cat. No. 339306, GeneGlobe ID. YP00203954), miR-148a-3p (QIAGEN, Hilden, NW, Germany, Cat. No. 339306, GeneGlobe ID. YP00205867), miR-223-3p (QIAGEN, Hilden, NW, Germany, Cat. No. 339306, GeneGlobe ID. YP00205986), miR-30a-5p (QIAGEN, Hilden, NW, Germany, Cat. No. 339306, GeneGlobe ID. YP02104140), and let-7b-5p (QIAGEN, Hilden, NW, Germany, Cat. No. 339306, GeneGlobe ID. YP00204750). We used the following thermal PCR cycle program: denaturation step at 95 □ for 2 min, followed by 40 cycles of denaturation at 95 □ for 10 s and annealing/elongation at 56 □ for 1 min. Note that samples collected from TIM-1 at T0 (before milk addition into the system) were also reverse transcribed, and their qPCR cycle of quantification (Cq) was considered as background. This background was subtracted from the Cqs of the samples collected at the other time points. Cq values of the spike-ins UniSp2 and UniSp6 were utilized to assess RNA extraction and RT-qPCR efficiency and for qPCR normalization (**Supplementary Fig.S2A-C**).

### Standard curve for absolute RNA quantification

The copy numbers of miR-30a-5p, miR-148a-3p, let-7b-5p, miR-223-3p were quantified using a standard curve generated with synthetic RNA oligonucleotides (IDT, Coralville, IA, USA; miR-148a-3p synthetic RNA sequence: 5’- UCAGUGCACUACAGAACUUUGU-3’; let-7b-5p synthetic RNA sequence: 5’- UGAGGUAGUAGGUUGUGUGGUU-3’; miR-30a-5p synthetic RNA sequence: 5’- UGUAAACAUCCUCGACUGGAAG-3’, miR-223-3p synthetic RNA sequence: 5’- UGUCAGUUUGUCAAAUACCCCA-3’). These oligonucleotides were serially diluted 1:10 to achieve concentrations ranging from 6×10 □ to 6×10^3^ copies (per qPCR well), spanning seven orders of magnitude. For each standard curve, the cycle of quantification (Cq) value was plotted against the corresponding copy numbers. A linear regression was performed to calculate the equation of the curve and the correlation coefficient (R^2^) (**Supplementary Fig.S3A-D**).

### Dye staining

A diluted stock of DiR (Thermo Fisher Scientific, Waltham, MA, USA, Cat. No. D12731) deep red dye, (Thermo Fisher Scientific, Waltham, MA, USA, Cat. No. C34565) or green dye (Thermo Fisher Scientific, Waltham, MA, USA, Cat. No. C7025) was prepared to a working concentration of 1.6 mM. In separate tubes, EVs were mixed or not with the diluted dye and then transferred to a clear 96-well plate. The plate was incubated at 37 °C for 30 min with gentle shaking. The reading was then performed using the VarioSkan plate reader (Thermo Fisher Scientific, Waltham, MA, USA).

### Statistical analysis

All statistical analyses were performed using Prism 10.4.1 (GraphPad Software Inc., San Diego, CA, USA). Given the small sample size (n=2), the normality test can not be applied. Thus, non-parametric tests like Mann-Whitney (for the comparison of two non-paired groups) and Kruskal-Wallis tests were used.

## Results

### HM microRNAs resist differentially to infant simulated digestion

To assess the stability of HM microRNAs against infant digestion, whole HM was digested in vitro using the TIM-1 system over two hours **(Figure 1A)**. The digestion parameters were set to simulate the digestion of infants aged one to six months **(Figure 1B)**. The absolute levels of the three microRNAs, miR-148a, let-7b, and miR-223, were assessed in each compartment of the system (stomach, duodenum, jejunum, and ileum) at 30 min and 2 h after the onset of digestion. The levels displayed correspond to the copy number in the total volume of each compartment at each time point. At T30 min, all three microRNAs were still detectable in the stomach **(Figure 1C)**. Surprisingly, their levels in the stomach were close to those in the total milk sample added to the system at T0 (input). In addition, both miR-148a and let-7b were detectable in the duodenum at T30 min, whereas miR-223 showed no detectable signal in that compartment. In the jejunum, miR-223 and let-7b were no longer detected, while miR-148a remained detectable at slightly lower levels than in the duodenum. None of the three microRNAs was detected in the ileum at this time point **(Figure 1C)**, as only 0.4% of the initial milk volume had reached this compartment. Notably, after 2 h of digestion (at T120 min), only miR-148a was still detectable in the duodenum, jejunum, and ileum **(Figure 1D)**, showing a marginal decrease in copy number compared to its levels at T30 min. Approximately 2.5 × 10□, 2.1 × 10□, and 3.8 × 10□ copies of miR-148a remained in the duodenum, jejunum, and ileum, respectively. In contrast, both let-7b and miR-223 were not detected in any compartment after 2 h digestion. Note that, at 2 h, the stomach was nearly empty, and sampling from this compartment was hence not possible. Effluent samples, representing the fraction of digesta that resisted upper gastrointestinal digestion and would reach the colon, were collected at different time points (T30, T60, T90, and T120 min). Notably, only miR-148a was detected in these samples, highlighting its stability throughout gastrointestinal transit (the copy number represents the sum of absolute levels found in effluents across all time points).

**Figure 1.**
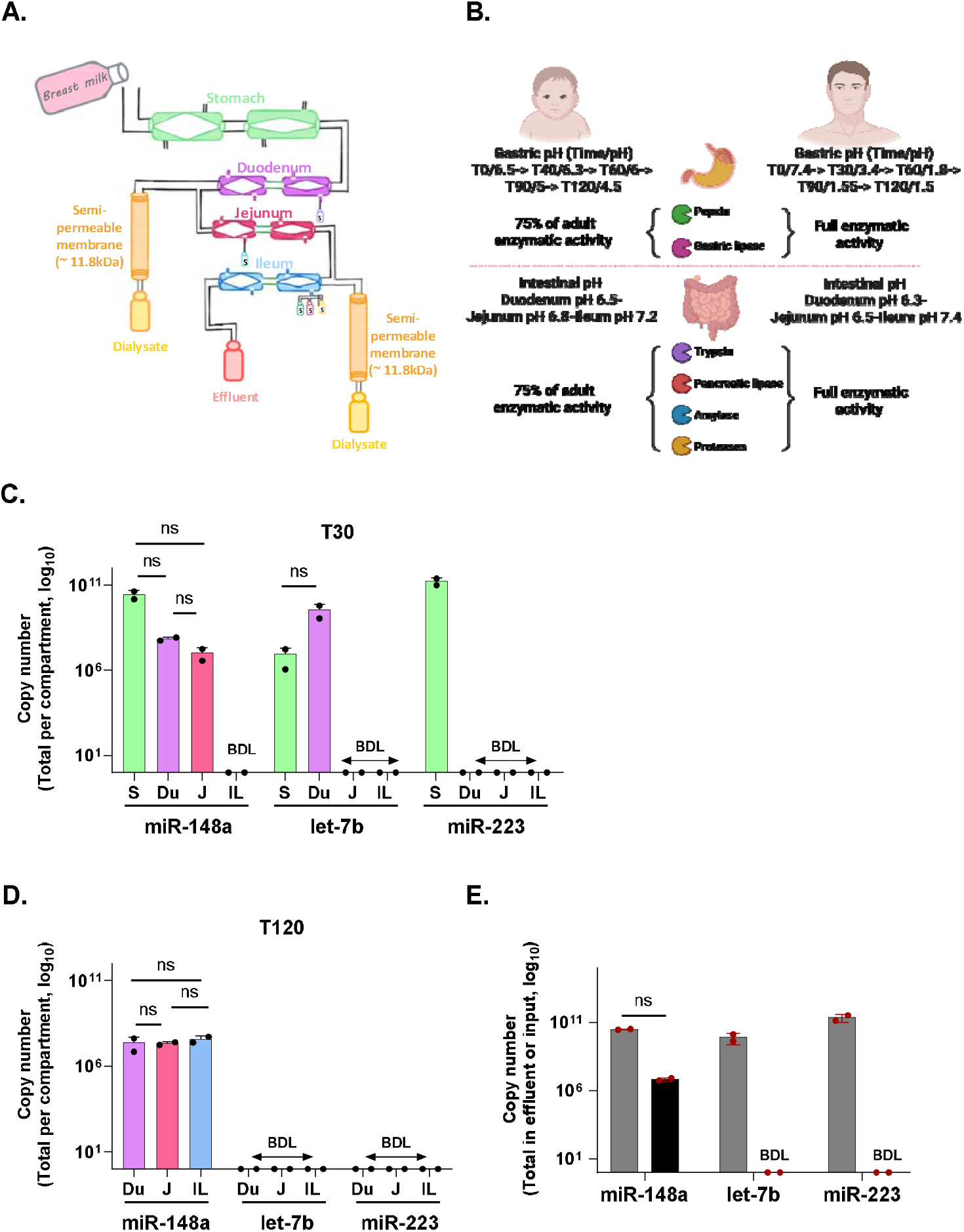
*In vitro* digestion of breast milk under conditions which simulate infant’s digestion. **A.** Schematic depicting the different compartments of the TNO TIM-1 *in vitro* digestion model. **B**. Schematic showing the modifications applied to the digestion parameters, compared to those of adults, to simulate an infant’s digestion. **C**. Quantification of three milk microRNAs (miR-148a, let-7b, and miR-223) in the different compartments of the TIM-1 model at T30 (after 30 min of digestion start) and **D**. at T120 (after 120 min of digestion). **E**. Copy number of the three milk microRNAs before (input milk added to the TIM-1 system) and after 2 h (120 min) of complete digestion. The effluent represents the digests that have passed through all system compartments and exited the system. Data represent the means of two independent digestions; one milk lot was used per digestion (n=2). The non-parametric Mann-Whitney test was used to assess statistical significance between input and effluent copy numbers of each microRNA. ns: non-significant, BDL: below detection limit. Signals below the detection limit were excluded from the statistical analysis.

### Characterization of EV populations isolated from HM

To check for the presence of the distinct EV populations, which were identified previously in commercial bovine milk ^(18)^, we subjected HM to differential ultracentrifugation and analyzed the components of the 12K, 35K, 70K, and 100K pellets obtained (**Figure 2A**). We first confirmed that the pellets were more enriched in vesicular components like lipid membrane and cytoplasmic proteins compared to the using the DiR dye and the green dye, respectively (**Supplementary Fig.S4A-B**). The DLS-based analysis of the size distribution of each pellet showed a single, relatively narrow, diameter peak, indicating a rather homogeneous population of EVs with a uniform size distribution, which is in accordance with the dispersity index values **(Figure 2B and Supplementary Fig.S4C)**. On average, the hydrodynamic diameter was 170 nm, 120 nm, 90 nm, and 70 nm for the 12K, 35K, 70K, and 100K populations, respectively **(Figure 2C)**. The hydrodynamic diameters obtained here for HM EVs were smaller than those previously obtained for EVs isolated from commercial bovine milk, as well as raw bovine milk ^(18)^. The values of the derived count rate (an estimate of particle count) in the 12K subset were the highest and tended to decrease progressively in the other three pellets **(Figure 2D)**. In accordance with the derived count rate, total protein content was highest in the pellets sedimented at low speed and decreased progressively in those sedimented at higher speeds **(Figure 2E)**. The majority of proteins did not co-precipitate with the pellets and remained in the SN **(Supplementary Fig.S4D)**. We then looked for five common milk EV protein markers (ALIX, TSG101, CD63, and Hsp70) known to play critical roles in multiple EV biogenesis pathways (Endosomal Complex Required for Transport (ESCRT)-dependent and ESCRT-independent) ^(44)^. Of note, TSG-101 showed two bands in all four pellets. The band detected at 75 kDa was not detected in previous data in either commercial or raw bovine milk EVs ^(18)^. We also detected the protein xanthine dehydrogenase (or XDH, which is known to associate with milk fat globules but was also found associated with milk sEVs (“exosomes”) ^(45)^) in all four pellets. Overall, the enrichment profile of the EV markers over the different EV subsets was similar to that observed in commercial bovine milk EVs ^(18)^.

**Figure 2.**
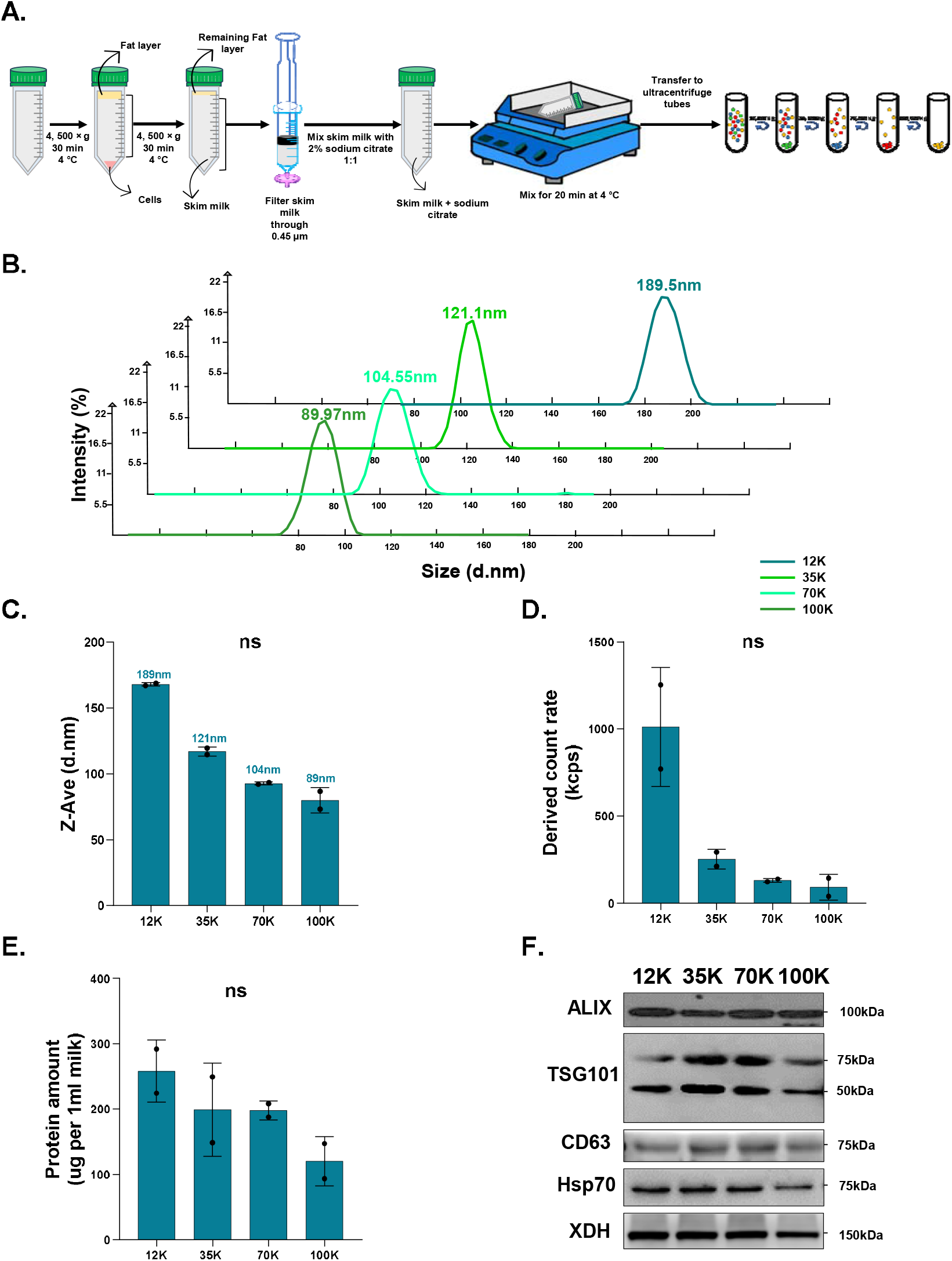
Isolation and characterization of human breast milk EV populations. **A.** Schematic depicting the different steps of human breast milk defatting and differential ultracentrifugation-based isolation of the four EV populations. **B**. Raw peaks showing the size distribution by intensity of particles in each of the four 12K, 35K, 70K, and 100K populations (presented are the averaged peaks of the two milk lots (biological duplicate) with three readings recorded per sample (technical triplicates)). **C**. Hydrodynamic diameter (Z-Ave, d.nm) of the four EV populations. Data are shown as mean ± SD (two milk lots, n=2). Statistical significance was assessed using the Kruskal–Wallis test followed by Dunn’s multiple comparisons test. **D**. DLS measurements of the particle count in the four EV populations. Data are shown as mean ± SD (two milk lots, n=2). Statistical significance was assessed using the Kruskal–Wallis test followed by Dunn’s multiple comparisons test. **E**. Quantification of the protein content (in μg) in 1 ml of milk EVs. The values are means ± SD of the two milk lots (n=2). The non-parametric Kruskal-Wallis test followed by Dunn’s multiple comparisons was used to assess statistical significance. **F**. Western blot of milk EV protein markers. ns: non-significant

### Highly abundant milk microRNAs are detected in the four HM EV populations

In many studies investigating the microRNA profile in human or bovine milk with an RNA sequencing approach, the same set of microRNAs, around ten to fifteen microRNAs, was reported to consistently dominate the detected reads ^(46)^. We chose to quantify three of these microRNAs, being miR-148a, let-7b and miR-30a, and study their distribution over the four EV populations in HM. The absolute levels (per 1 ml milk) of each microRNA indicated that, among the three microRNAs, let-7b appeared to be the most highly enriched with up to 5.2 × 10^7^ copies per 1 ml milk. As reported before, miR-148a came directly after let-7b-5p with up to 1.4 × 10^6^ copies per 1 ml milk, while miR-30a was at around 1.3 × 10^6^ copies per 1 ml **(Figure 3A)**. As was shown in previous reports ^(18,47)^, the three microRNAs investigated were majorly enriched in the EV fraction compared to milk serum, which is represented here by the supernatant obtained after the last centrifugation at 100,000 × g. More than 90% of total microRNA levels were detected in all four EV fractions, while less than 10% were detected in the SN **(Figure 3B)**. All three microRNAs showed similar distribution over the four EV populations, where they were all mostly enriched in the 12K fraction, and their levels tended to decrease in the lighter EV populations (35K, 70K, and 100K) **(Figures 3C, D and E)**.

**Figure 3.**
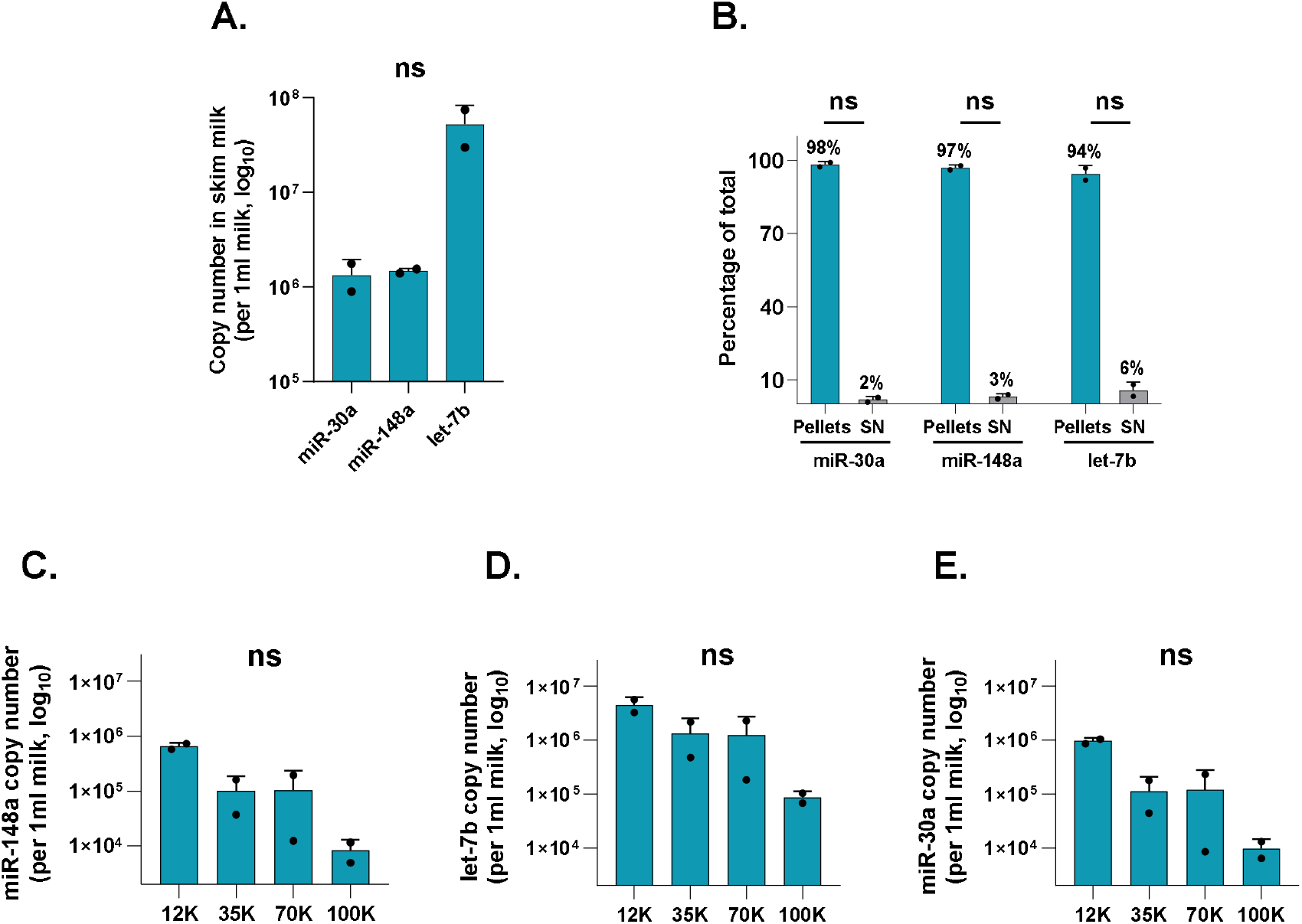
Absolute quantification of three milk microRNAs in human breast milk and milk-derived EV samples. **A.** Absolute quantification of the three milk microRNAs, miR-148a, let-7b, and miR-30a, in non-fractioned human breast skim milk. Data are shown as mean ± SD (two milk lots, n=2). Statistical significance was assessed using the Kruskal–Wallis test followed by Dunn’s test for multiple comparisons between the different microRNAs. **B**. Partitioning of the three microRNAs over the EV populations and the supernatant (SN, or milk serum; obtained from the last centrifigation step). Data are shown as mean ± SD (two milk lots, n=2). The non-parametric Mann-Whitney test was used to perform pairwise comparisons between SN and pellets for each microRNA. Absolute quantification of the three milk microRNAs **C**. miR-148a, **D**. let-7b, and **E**. miR-30a in the four EV populations. Copy number is presented per 1 ml of milk EVs. Data are shown as mean ± SD (two milk lots, n=2). Statistical significance was assessed using the Kruskal–Wallis test followed by Dunn’s test for multiple comparisons between EV populations. Data are shown as mean ± SD (two milk lots, n=2). The exogenous synthetic UniSp2 spike-in was used as a qPCR normalizer to control for variations in RNA isolation and RT-qPCR efficiency. ns: non-significant

## Discussion

The potential transfer of functionally active dietary microRNAs, including milk microRNAs, is widely debated due to uncertainties about their stability during digestion and their ability to reach target sites ^(48)^. Compared with adults, infants have milder digestive conditions and a more permeable gastrointestinal (GI) barrier, which may favor the survival of microRNAs. In this study, we show that three highly abundant and predominant microRNAs from pasteurized HM exhibit differential resistance to infant GI digestion: while let-7b, miR-148a, and miR-223 all partially resisted gastric digestion, only miR-148a survived the complete digestive process. Overall, microRNA survival appears to be microRNA-specific and select milk-derived microRNAs may persist in detectable forms, although their functional relevance remains to be established.

The extreme loss of miR-148a observed here contrasts with previous reports, highlighting the potential influence of bile and dynamic digestion conditions. Only 0.2 □ % of miR-148a was detectable after 2 h (considering both the effluent and the amounts remaining within the system compartments), a value markedly lower than that reported in earlier studies. In simulations of infant gastric and intestinal digestion in which samples were mixed with digestive enzymes under defined pH and electrolyte conditions, microRNAs contained in preterm raw HM exosomes showed minimal loss, not exceeding 9□ % in one study ^(32)^, while a subsequent report found overall exosomal microRNA levels to remain largely unchanged, with only a small subset exhibiting losses of up to 30□ % ^(33)^. To our knowledge, the present study is the first to incorporate bile into a dynamic infant digestion model, which likely accounts for much of the observed discrepancy. Additional factors, including differences in digestion conditions, duration, milk processing, and microRNA identity, may also contribute. Notably, pasteurized HM was used here, whereas previous studies analyzed exosomes from raw HM; pasteurization may compromise EV integrity and thereby increase microRNA susceptibility to degradation ^(34)^. Moreover, only three microRNAs were analyzed in the present study, and their differential stability further highlights that survivability may vary substantially across milk microRNA species.

In line with our findings, studies simulating adult digestion further support a key role for bile and intestinal conditions in the extensive degradation of microRNAs. A study assessing miR-148a transfected into raw bovine milk EVs and exposed to digestive enzymes and bile salts under defined pH and electrolyte conditions reported very low stability (0.26□ % remaining after 3□ h), with greater loss during the intestinal than the gastric phase ^(49)^. Using the TIM-1 system, another study measured two milk microRNAs (bta-miR-223 and bta-miR-125b) after adult digestion and reported 12□ % and 26□ % remaining after 2□ h, respectively ^(25)^. Although quantitatively higher than the 0.2□ % observed in pasteurized HM, these values reflect the same overall trend of pronounced microRNA loss. Mechanistically, bile acids can destabilize lipid membranes, disrupt EVs, and expose their cargo ^(50)^, while infant-like gastric pH may reduce protein aggregation and matrix-mediated protection, rendering EVs more susceptible to degradation. Consistent with this, studies on saliva and milk EVs have shown that bile in combination with intestinal enzymes can rupture EV membranes and degrade vesicular proteins ^(51,52)^. Furthermore, differences in milk composition, such as the higher casein content of bovine milk, compared to HM, may further influence microRNA stability ^(53)^. Overall, these observations indicate that digestive conditions and milk processing are important determinants of microRNA survival and help explain differences observed across digestion models.

Beyond survival, another important aspect of this study is the characterization of distinct EV populations in HM, similar to what has been reported in commercial bovine milk ^(18)^. We found that EVs sedimenting at low centrifugal forces harbor the majority of milk microRNAs. Such EVs have also been described in other biological fluids and media ^(54,55)^, but they are often discarded in standard EV isolation procedures or co-isolated with smaller EVs as heterogeneous populations ^(56–58)^. To better resolve this complexity, we adapted the protocol developed by Benmoussa et al. ^(18,39)^ to selectively isolate and characterize different EV populations from pasteurized HM. These EV populations differed in physical properties, including size and particle number, as well as in microRNA content. Although proteomic and lipidomic profiling were not performed in this study, previous work in bovine milk has shown that comparable EV populations possess distinct proteomic signatures ^(19)^. Such compositional differences may influence cellular uptake pathways and the fate of EV cargo, including microRNAs ^(59)^, and may also affect resistance to digestion and physiological effects in consumers ^(11,25,60)^. For example, in a colitis model, bovine milk EVs isolated at 100K and 35K exerted distinct biological effects: 100K EVs influenced intestinal microRNA levels and ZO-1 expression, whereas 35K EVs modulated innate immune responses ^(11)^.

Overall, differences in digestion models, digestion conditions, milk processing, and the microRNA species analyzed across studies complicate direct comparisons of milk microRNA stability during digestion. Nevertheless, most studies consistently report the presence of residual microRNAs after digestion, raising the question of whether the remaining amounts are sufficient to exert canonical microRNA functions. This question is particularly relevant given reports describing the detection of orally administered bovine milk EVs and microRNAs in mouse tissues ^(61–63)^. In this context, the present study provides the first assessment of microRNA stability from pasteurized HM under infant dynamic digestion conditions and explores the different EV populations in pasteurized HM. However, the study is limited by the small number of milk samples analyzed (two pooled lots, each representing 36 donor mothers) and by the lack of pre-pasteurization material, which precluded a direct evaluation of the impact of pasteurization on microRNA abundance.

## Conclusion

In conclusion, our results indicate that a substantial fraction of milk microRNAs is lost during infant digestion, with only select microRNAs, such as miR-148a, remaining detectable. This loss is likely influenced by dynamic digestive conditions, bile, and milk pasteurization. These findings provide important context for interpreting microRNA bioaccessibility and potential bioactivity in human milk, highlighting that microRNA content and stability may differ between raw donor milk, pasteurized milk, and infant formulas. Given the potential role of microRNAs in infant development, the impact of pasteurization on bioactive milk components in human milk banks warrants further evaluation. This raises the possibility that similar bioactive components, including microRNAs, may also be compromised or absent in infant formulas.

## Author contributions

Z.H. conceived and designed the study. Z.H. and E.P. adapted and performed the TIM-1 experiments. Z.H. performed the rest of the experiments and collected the data. Z.H. analyzed the data. All authors read and approved the final manuscript.

## Acknowledgements

We thank Dr. Patrick Provost for his major role in this project, including designing the study, supervising the project, designing key experiments, and providing the funding support. This work was supported by the Canadian Institutes of Health Research (CIHR) (SIRUL: 123293, to Dr. Patrick Provost). We also thank Dr. Mélissa Girard for coordinating the collection of milk samples and overseeing their transfer from Héma-Québec. We also extend our sincere thanks to Dr. Caroline Gilbert for the invaluable guidance in supervising the writing process of this paper and for her assistance with the figures and data analysis. We acknowledge the use of artificial intelligence– assisted tools, including Copilot (https://www.microsoft.com/en-us/microsoft-copilot) and ChatGPT (OpenAI; https://openai.com/chatgpt), to improve sentence structure and to look for the most relevant literature. All final decisions regarding content, interpretation, and citations were made by the authors.

## Data availability statement

No new datasets were generated.

## Abbreviations

HM: Human milk
EVs: Extracellular vesicles
IGF-I: Insulin-like growth factor-I
IGF-II: Insulin-like growth factor II
EGF: Epidermal growth factor
GI: Gastro-intestinal
DLS: Dynamic light scattering
Cq: Cycle of quantification
SDS: Sodium dodecyl sulfate
TBS: Tris buffer saline
SN: Supernatant
ESCRT: Endosomal complex required for transport
ZO-1: Zonula Occludens-1.

