## Supplemental figures for "Human milk contains a heterogeneous population of EVs and microRNAs that resist simulated digestion"

**Supplementary data**

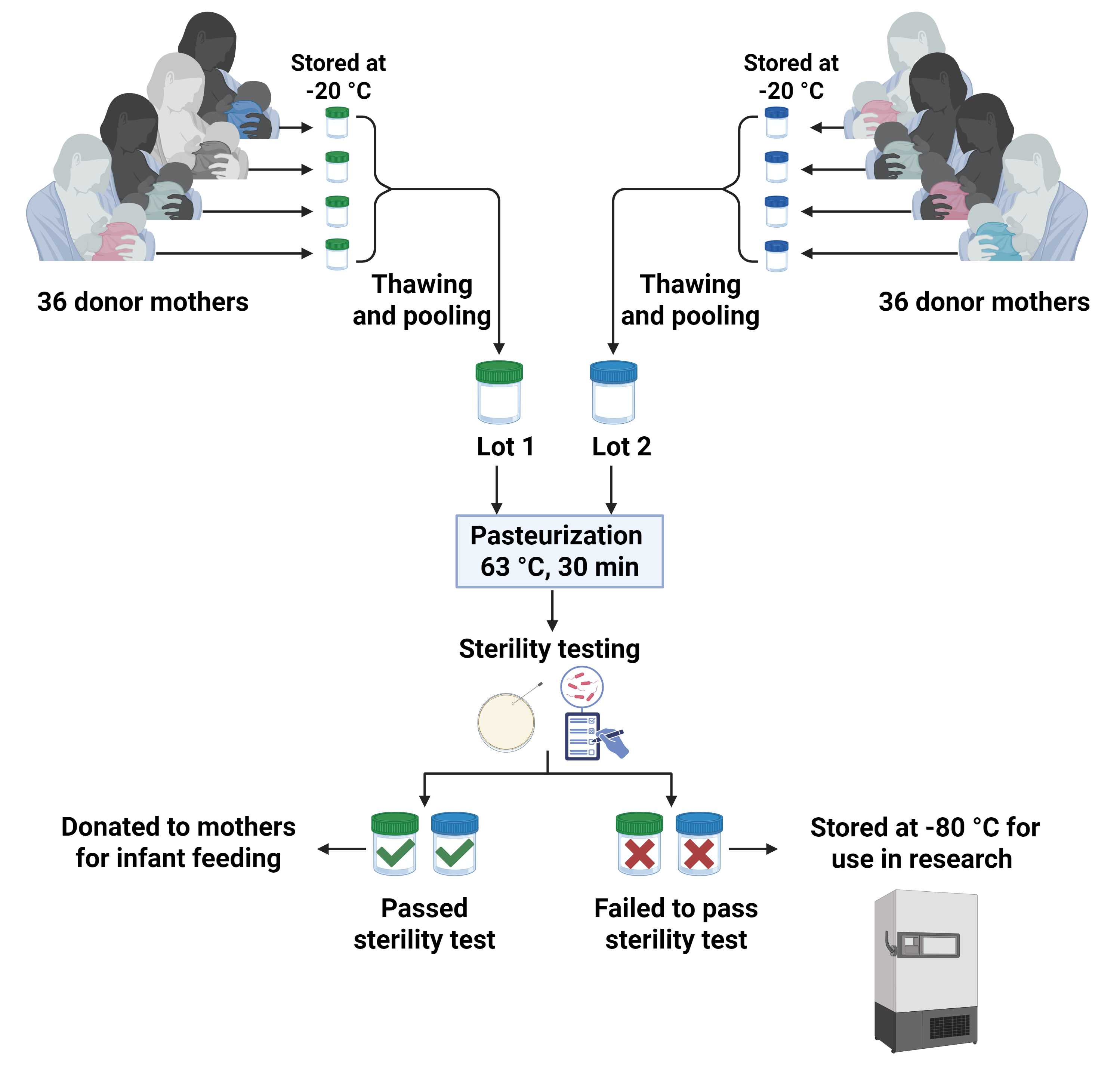

**Supplementary Fig.S1.**

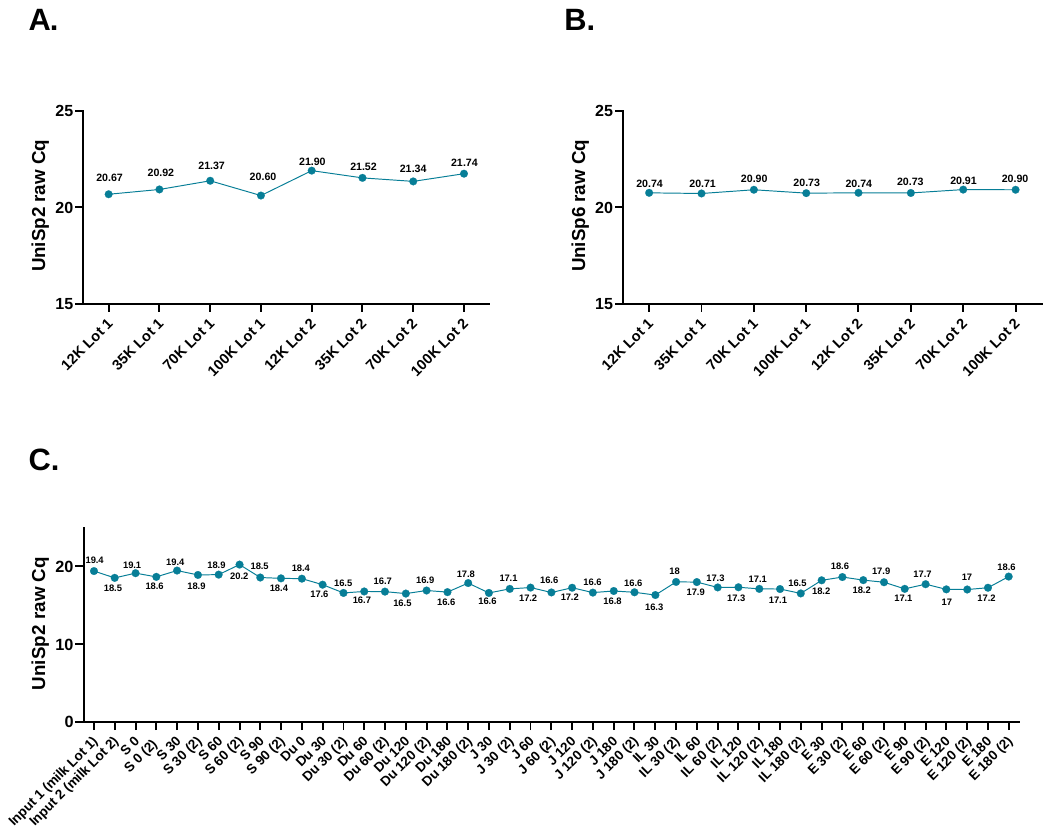

**Supplementary Fig.S2.**

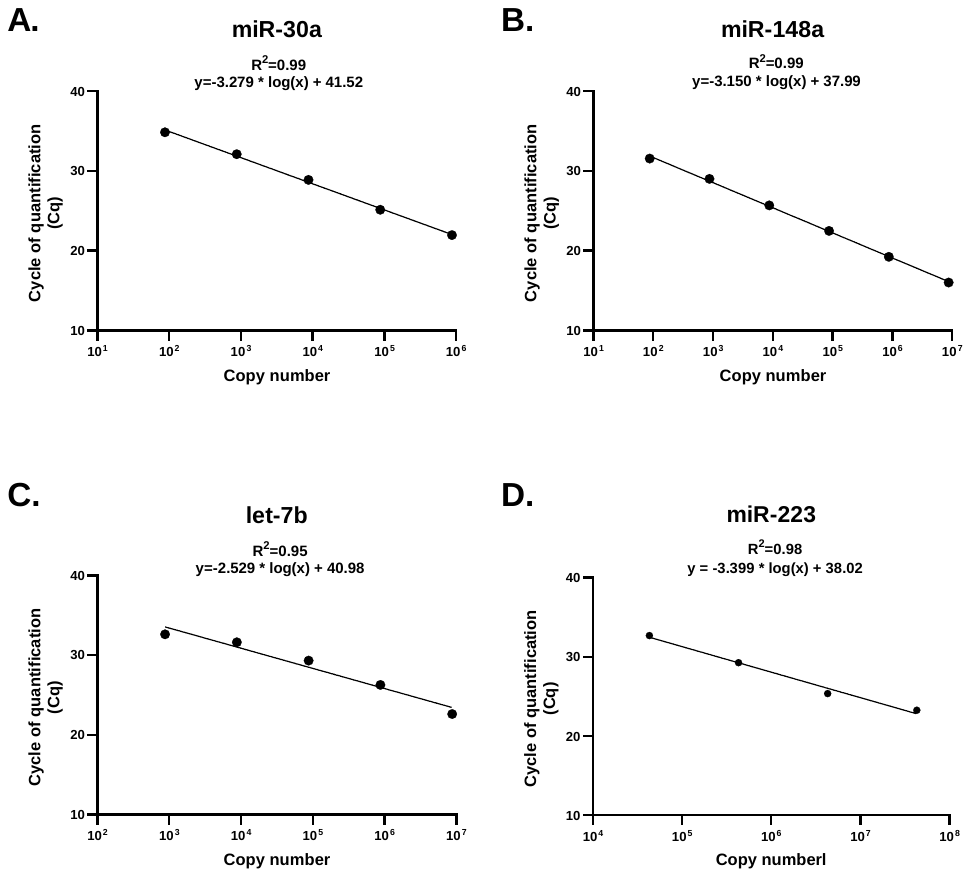

**Supplementary Fig.S3.**

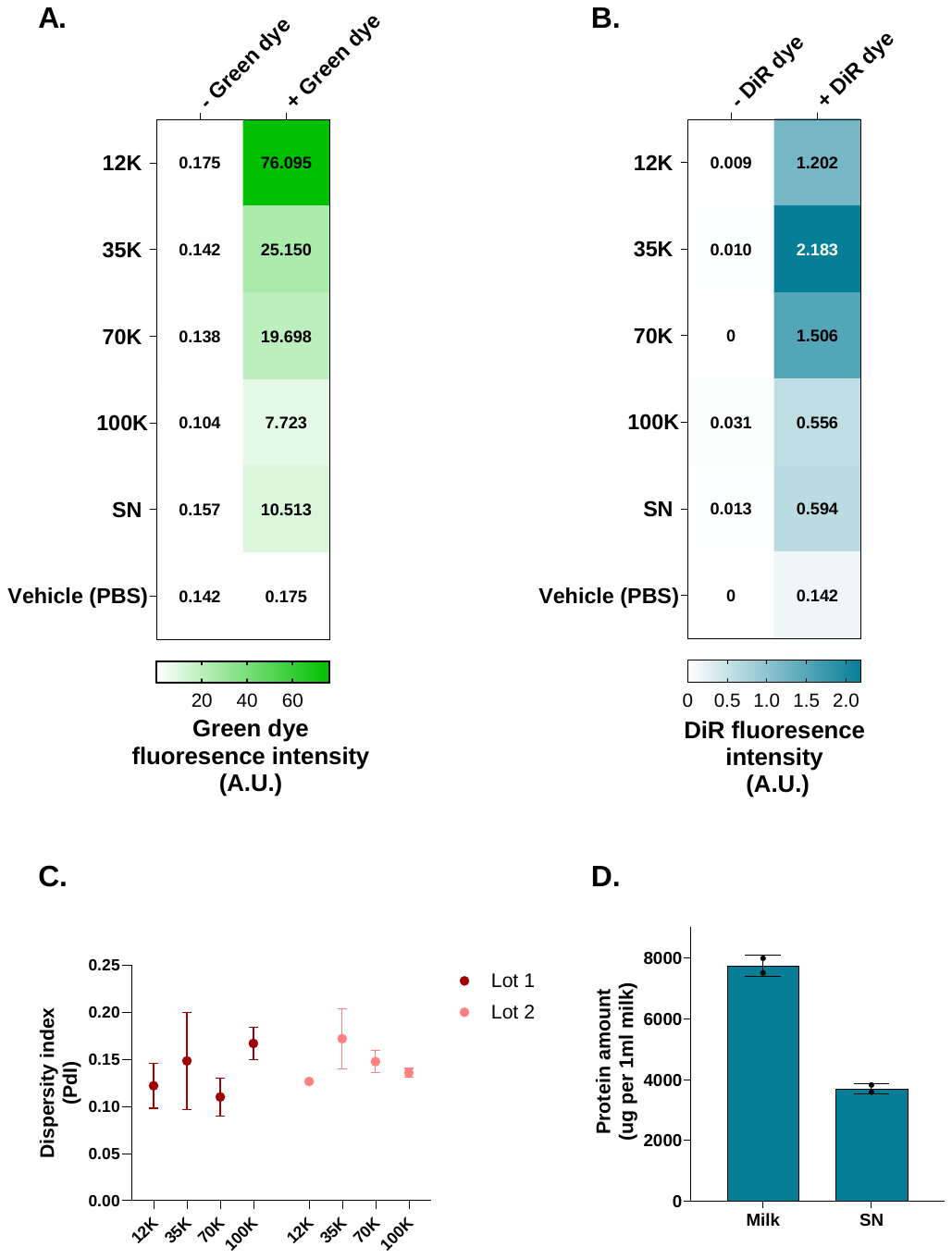

**Supplementary Fig. S4**

**Supplementary figure legends**

**Supplementary Fig.S1.** **Collection of human breast milk samples.** Human milk samples used in this study were obtained from the Public Mothers’ Milk Bank at Héma-Québec center, Quebec City, Canada. Two milk lots were used in the current study. Each milk lot (1 liter) was prepared by pooling samples from 36 donor mothers. The lots were then pasteurized at 63 °C for 30 min and subsequently tested for sterility. Samples that fail to pass the sterility test are deemed unfit for infant consumption, kept at – 80 °C, and repurposed for research. Samples were transferred from the milk bank to our laboratory on dry ice and promptly stored at −80 °C while still frozen, pending further analysis.

**Supplementary Fig.S2. RNA extraction and RT-qPCR quality control.** Raw cq values of the two exogenous spike-in RNA molecules used to control and normalize for variations during RNA extraction and RT-qPCR. **A.** UniSp2 and **B.** UniSp6 for EV samples, and **C.** UniSp2 for TIM-1 samples.

**Supplementary Fig.S3. Standard curves for microRNA RT-qPCR-based absolute quantification.** Standard curves showing the copy number (in qPCR well) of **A.** miR-30, **B.** miR-148a, **C.** let-7b, and **D.** miR-223 were established using the synthetic RNA oligonucleotides (IDT, Coralville, IA, USA) serially diluted 1/10th to obtain concentrations of 4-6 logs.

**Supplementary Fig.S4. Characterization of human breast milk EV populations.** Heat maps showing the fluorescence intensities of the **A.** Green dye and **B.** DiR lipophilic dye. Milk EV populations were labeled with the aforementioned dyes to confirm the presence of vesicular components (e.g., lipid bilayer membrane and cytosolic proteins). The Green dye can traverse the lipid bilayer of cells as well as other lipid-bilayered components, where it fluoresces upon catalysis by the cytosolic glutathione-S-transferase. The DiR dye penetrates through the membrane and becomes fluorescent after integrating into the lipid components of the bilayer membrane. **C.** Dispersity index measured by DLS as an estimate of the size uniformity of particles in the different EV populations of each milk lot. Reported errors are the standard deviation of three measurements. **D.** Quantification of the protein content (in ug) in 1 ml of SN or skim milk. The non-parametric Mann-Whitney test was used for pairwise comparisons of protein amounts (n=2).
